# Heterozygous Inversion Breakpoints Suppress Meiotic Crossovers by Altering Recombination Repair Outcomes

**DOI:** 10.1101/2022.11.09.515852

**Authors:** Haosheng Li, Erica Berent, Savannah Hadjipanteli, Miranda Galey, Danny E. Miller, Nicole Crown

## Abstract

Heterozygous chromosome inversions suppress meiotic crossover (CO) formation within an inversion, potentially because they lead to gross chromosome rearrangements that produce inviable gametes. Interestingly, COs are also severely reduced in regions nearby but outside of inversion breakpoints even though COs in these regions do not result in rearrangements. Our mechanistic understanding of why COs are suppressed outside of inversion breakpoints is limited by a lack of data on the frequency of noncrossover gene conversions (NCOGCs) in these regions. To address this critical gap, we mapped the location and frequency of rare CO and NCOGC events that occurred outside of the *dl-49 chrX* inversion in *D. melanogaster*. We created full-sibling wildtype and inversion stocks and recovered COs and NCOGCs in the syntenic regions of both stocks, allowing us to directly compare rates and distributions of recombination events. We show that COs are completely suppressed within 500 kb of inversion breakpoints, are severely reduced within 2 Mb of an inversion breakpoint, and increase above wildtype levels 2-4 Mb from the breakpoint. We find that NCOGCs occur evenly throughout the chromosome and, importantly, occur at wildtype levels near inversion breakpoints. We propose a model in which COs are suppressed by inversion breakpoints in a distance-dependent manner through mechanisms that influence DNA double-strand break repair outcome but not double-strand break location or frequency. We suggest that subtle changes in the synaptonemal complex and chromosome pairing might lead to unstable interhomolog interactions during recombination that permits NCOGC formation but not CO formation.

## Introduction

Chromosome inversions have far-ranging impacts on reproduction and speciation when paired with a non-inverted homolog. At the molecular level, heterozygous inversions disrupt fundamental aspects of meiosis by suppressing both the formation and recovery of meiotic crossovers (COs) within the inversion and in the regions nearby but outside the inversion breakpoints (1). At the population level, suppressing COs prevents genetic exchange between an inversion and its non-inverted homolog, dramatically reducing gene flow for that portion of the genome (2, 3). Within the inversion, recombination is unable to separate combinations of alleles, which can be beneficial for advantageous or adaptive alleles (4–6). Alternatively, suppression of exchange can have negative outcomes such as harboring selfish genetic elements and meiotic drive systems (reviewed in (7)). Lastly, the ability of inversions to prevent gene flow locally in the genome is the basis of the chromosomal theory of speciation (8, 9). Given the far-ranging impacts of heterozygous inversions on reproduction and speciation, it is critical to understand how they disrupt meiosis at the molecular level. Despite a rich history of studying inversions, the mechanisms of how they suppress COs remains unknown. Here, we exploit the unique properties of inversion breakpoints to provide crucial insight into the mechanisms of CO suppression in *Drosophila melanogaster*.

Early work on heterozygous inversions focused on two possible explanations for how they might suppress COs. The first possibility is that when inversions are heterozygous with a non-inverted chromosome, meiotic chromosome pairing and synapsis are defective, which prevents CO formation (10, 11). The second possibility is that COs can form, but because COs within the inversion will lead to chromosome rearrangements, the gametes containing such chromosomes will be inviable (1, 11) (Figure 1). This will make it appear as if COs are suppressed, when in reality, those chromosomes simply are not recovered in the offspring. This debate was mostly settled in 1936 (12, 13) when Sturtevant demonstrated that COs do form within an inversion yet, paradoxically, there is no increased embryonic lethality as expected; he suggested a model where chromosomes resulting from COs within an inversion are aligned on the meiotic spindle such that they are preferentially placed in the polar bodies and eliminated (see (14) for a summary of this model). Despite the lack of direct cytological evidence for Sturtevant’s model, the field has generally accepted the notion that heterozygous inversions suppress COs because the aneuploid offspring are not recovered (15–17).

**Figure 1.**
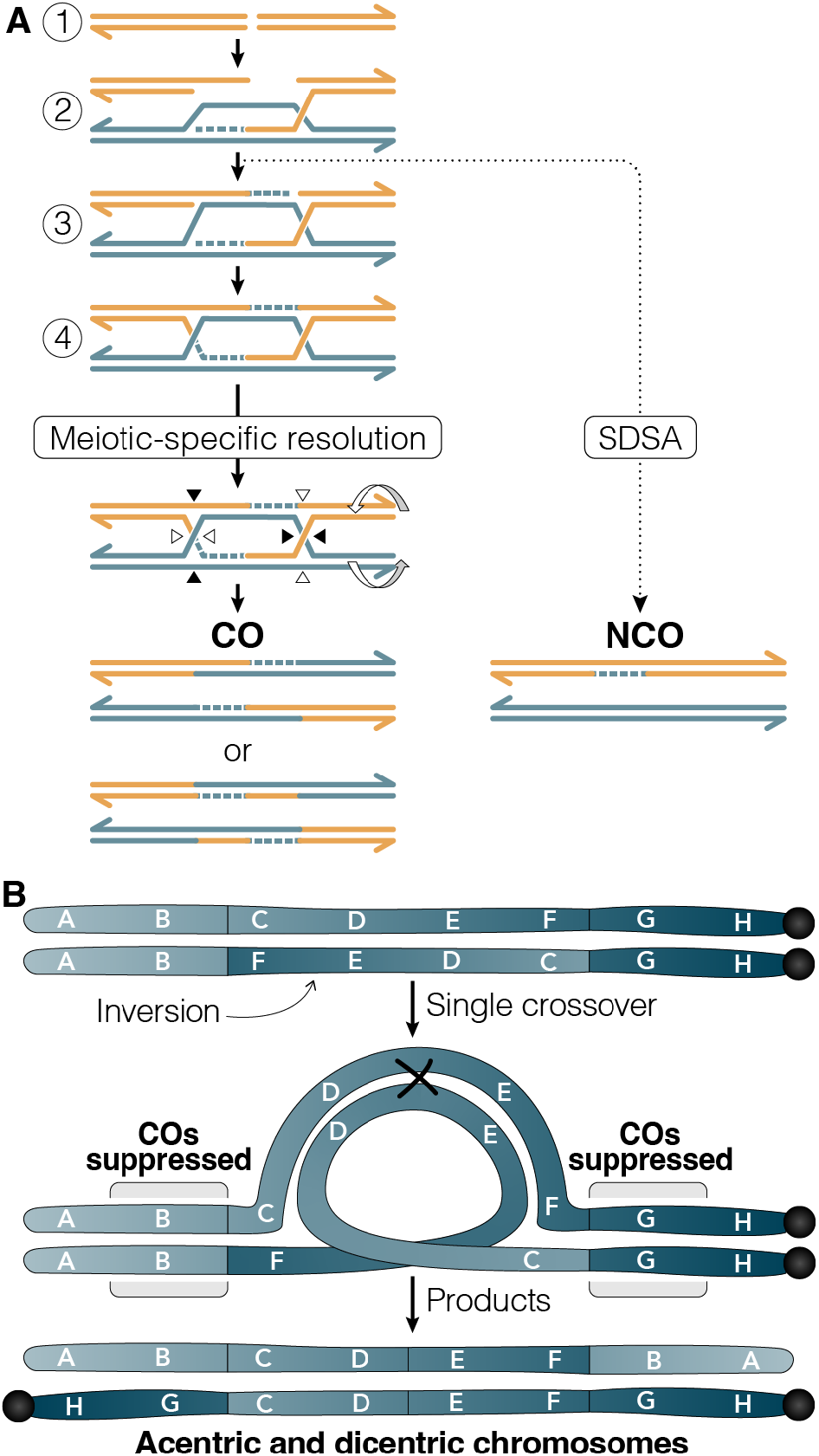
A) Canonical recombination pathways used in meiosis. 1) Meiosis is initiated by an enzymatically mediated DSB. 2) The DSB is resected into single-stranded ends, one of which invades the homologous chromosome and primes DNA synthesis. This forms a displacement loop (D-loop) and during synthesis dependent strand annealing (SDSA), this structure can be unwound by a helicase into a NCO. 3) If the second end of the DSB is captured by the D-loop, it also primes synthesis. 4) The second-end capture intermediate is ligated into a double Holliday Junction, which is cleaved by a meiosis-specific endonuclease to form exclusively COs. B) Predicted pairing arrangement between an inversion and a standard arrangement homologous chromosome. Single COs within the inversion breakpoints lead to acentric and dicentric chromosomes with deletions and duplications. COs are also suppressed in the collinear regions outside of the inversion breakpoints, even though COs in these regions would not lead to chromosome rearrangements.

Concomitant with the early observation that COs are suppressed within heterozygous inversions was the observation that they are also severely reduced in regions immediately outside the inversion breakpoints (11, 12, 18). For example, on a chromosome carrying the *dl-49* inversion in *D. melanogaster*, COs are reduced to approximately 30% of wildtype in the proximal interval and to about 4% of wildtype in the distal interval (11). This phenomenon is not exclusive to *D. melanogaster* as it was also seen in *D. pseudoobscura* by Dobzhansky and Epling (4). Additionally, in interspecies crosses between *D. pseudoobscura* and *D. persimilis*, crossover frequency ranges from 0.0% to 0.5% in regions up to approximately 2.5 Mb outside of the inversion breakpoints, depending on the inversion (19, 20). Since COs that occur outside of the inversion breakpoints will not lead to chromosome rearrangements, there must be a mechanistic explanation for why they do not form in these regions.

COs are made by repairing a DNA double-stranded break (DSB) made during meiosis (Figure 1). However, only a few select DSBs will be repaired into COs, while the majority of DSBs will be repaired into noncrossovers (reviewed in (21)). If these noncrossovers are associated with gene conversions, then there are small tracts of unidirectional gene transfer from one homolog to the other. A major open question is if heterozygous inversions impact noncrossover gene conversions (NCOGCs) in a manner similar to crossovers.

Data are sparse on how heterozygous inversions affect NCOGCs because they are notoriously difficult to detect due to their small size, less than 1 kb in Drosophila species (22–24). It is possible to select for NCOGCs using purine selection for intragenic recombination events at the *rosy* locus in *D. melanogaster*, but these events are so rare that it requires selecting against 500,000 to 1 million offspring in each experiment (23). Previous work using this approach to study NCOGC frequencies in the interior of an inversion selected against 5,000,000 offspring to recover 66 NCOGCs; this work demonstrated that NCOGCs occur within a heterozygous inversion at the same frequency as in standard arrangement chromosomes (25). Even attempts at comprehensively assessing the frequency and location of NCOGCs using genome-wide analyses are limited by small sample sizes. Sequencing-based analyses in *D. pseudoobscura* and *D. persimilis* showed that NCOGC frequencies inside inversions were at least as high as in collinear regions, but the number of unique NCOGC events was only 32 (26). Similar sequencing analyses in *D. melanogaster* show that NCOGCs within inversions occur at rates higher than on standard arrangement chromosomes (27). While this work was able to analyze the location of 79 unique NCOGCs, it was done using multiply inverted balancer chromosomes, which may not be representative of how single or naturally occurring inversions behave.

The same sequencing-based analyses described above provide some limited insight into how NCOGCs behave outside of inversion breakpoints. In *D. melanogaster*, 20 of the 79 unique NCOGCs occurred between 0.5 and 1 Mb away from the inversion breakpoints and one NCOGC was detected only 14kb away (27). In the *D. pseudoobscura* and *D. persimilis* analysis, there was one NCOGC 37 kb away from an inversion breakpoint (26). The low — but non-zero — frequency of NCOGCs within 500 kb of inversion breakpoints suggests that these regions are at least competent to form DSBs, although it remains to be seen if DSBs form at the same frequency in these regions as in collinear portions of the genome. Critically, the datasets on NCOGCs outside inversion breakpoints are too small to draw robust mechanistic conclusions.

The emerging picture of recombination outside of inversion breakpoints is that CO frequency is severely reduced and that NCOGCs can occur. However, it remains unclear whether NCOGCs are reduced by similar levels as COs, preventing crucial insight into whether CO suppression is mediated by a reduction in DSBs or whether recombination is biased away from CO repair (Figure 1). To address this, we generated a high-resolution map of rare CO and NCOGC events outside of a single X chromosome inversion in *D. melanogaster*. Critically, we built full-sibling wildtype and inversion stocks and recovered recombination events in syntenic regions, enabling us to directly compare frequencies and distributions of recombination events. We find that COs are suppressed in a distance-dependent manner from the inversion breakpoint and that NCOGCs occur at wildtype frequencies outside of inversion breakpoints, suggesting that inversion breakpoints suppress COs by altering recombination outcomes as opposed to suppressing DSB formation.

## Methods

### Drosophila husbandry and stocks used

Flies were maintained at 25 °C on standard cornmeal media. The X chromosome inversion *dl-49, v^Of^, f^1^* was ordered from the Bloomington Drosophila Stock Center (stock #779). Previous sequencing of the *FM7* balancer chromosome showed the *dl-49* inversion breakpoints occur at nucleotide position 4,897,260 and between nucleotides 13,426,854 and 13,427,212 (28). We confirmed that these same inversion breakpoints are in the single *dl-49* inversion stock using PCR (Supplemental Figure 1) and Nanopore sequencing (data not shown). We isogenized *chrX* and *chr2* of the *dl-49* stock by first making a *dl-49, v^Of^, f^1^; Pin/CyO* stock. This stock was then crossed to a fully isogenized Oregon-RM stock (courtesy of R. Scott Hawley), and a single male of the genotype *dl-49, v^Of^, f^1^; +/CyO* was crossed back to *dl-49, v^Of^, f^1^; Pin/CyO* females. The male and female offspring of this cross that were *dl-49, v^Of^, f^1^; +/CyO* were crossed together to establish a stock that was *dl-49, v^Of^, f^1^; +/+*, with *chr2* in this stock descending from Oregon-RM. The *y^1^ cv^1^ IP3K2^wy-74i^ f^1^* stock was generated by isolating a recombinant between *cv^1^* and *IP3K2^wy-74i^* from *y^1^ cv^1^ v^1^ f^1^/sc^1^ ec^1^ IP3K2^wy-74i^ f^1^* females (Bloomington Drosophila Stock Center #1274). This new *y^1^ cv^1^ IP3K2^wy-74i^ f^1^* stock was partially isogenized by crossing the single male recombinant to *FM7w* females (29). We refer to *IP3K2^wy-74i^* as *wy* in the main text for clarity. The X and 2^nd^ chromosomes of the *y^1^w^1118^* stock were partially isogenized by crossing a single male to *FM7w; Pin/CyO* females. All stocks are available upon request.

### Generating full-sibling dl-49 and wildtype stocks

To control for differences in genetic background that might influence recombination rates (30), we generated full-sibling *dl-49* and Oregon-RM stocks. A fully annotated crossing scheme can be found in Supplemental Figure 2. Briefly, in generation 1, males from the previously isogenized *dl-49, v^Of^, f^1^; +/+* stock were crossed to previously isogenized Oregon-RM females (stock courtesy of R. Scott Hawley). In generation 2, heterozygous *+/dl-49, v^Of^, f^1^* females were crossed to sibling *Oregon-RM* males. In generation 3, a single male that contained the *dl-49, v^Of^* but had lost *f^1^* was crossed to homozygous *Oregon-RM* sisters. The goal of crossing off *f^1^was* to ensure at least one recombination event near the *dl-49* inversion breakpoint, which would increase the amount of shared genetic material between *dl-49* and the wildtype full-sibling stock. In generations four through nine, the *dl-49, v^Of^* chromosome was maintained in females in a heterozygous state to allow recombination between it and the wildtype *Oregon-RM* X chromosome. Additionally, in all nine generations, *chr2* and *chr3* were freely recombining. At the tenth generation, homozygous *dl-49, v^Of^* and wildtype *Oregon-RM* full-sibling stocks were established. It’s important to note that the *dl-49* chromosome, *chrX* and *chr2* are fully isogenized, while *chr3* has shared genetic background but is not isogenized.

### Recovering recombinants

Crossovers were recovered in two separate crosses, with slightly different experimental set ups (Figure 2). In the first experimental set up, we generated females that were *dl-49, v^Of^, f^1^/y^1^w^1118^*. These females were crossed to males of the reference genome stock *y; cn bw sp/CyO*. We collected individual male offspring that came from a CO between *y* and *f* in the maternal genome.

After completing our first experimental set up, we decided to control for genetic background by generating full sibling stocks as described above. To collect COs in the new full sibling stocks, we generated females that were *dl-49, v^Of^/y^1^ cv^1^ IP3K2^wy-74i^ f^1^* (Figure 2). These females were crossed to males of the reference genome stock *y; cn bw sp/CyO*. We again collected male offspring that came from a CO between *y* and *f* in the maternal genome. An unforeseen complication arose in this cross because we were unable to detect the difference between nondisjunction and a CO until the CO offspring were sequenced. While this prevents our ability to accurately calculate overall CO frequency in the interval, it does not affect our analysis of CO distributions within the interval.

**Figure 2.**
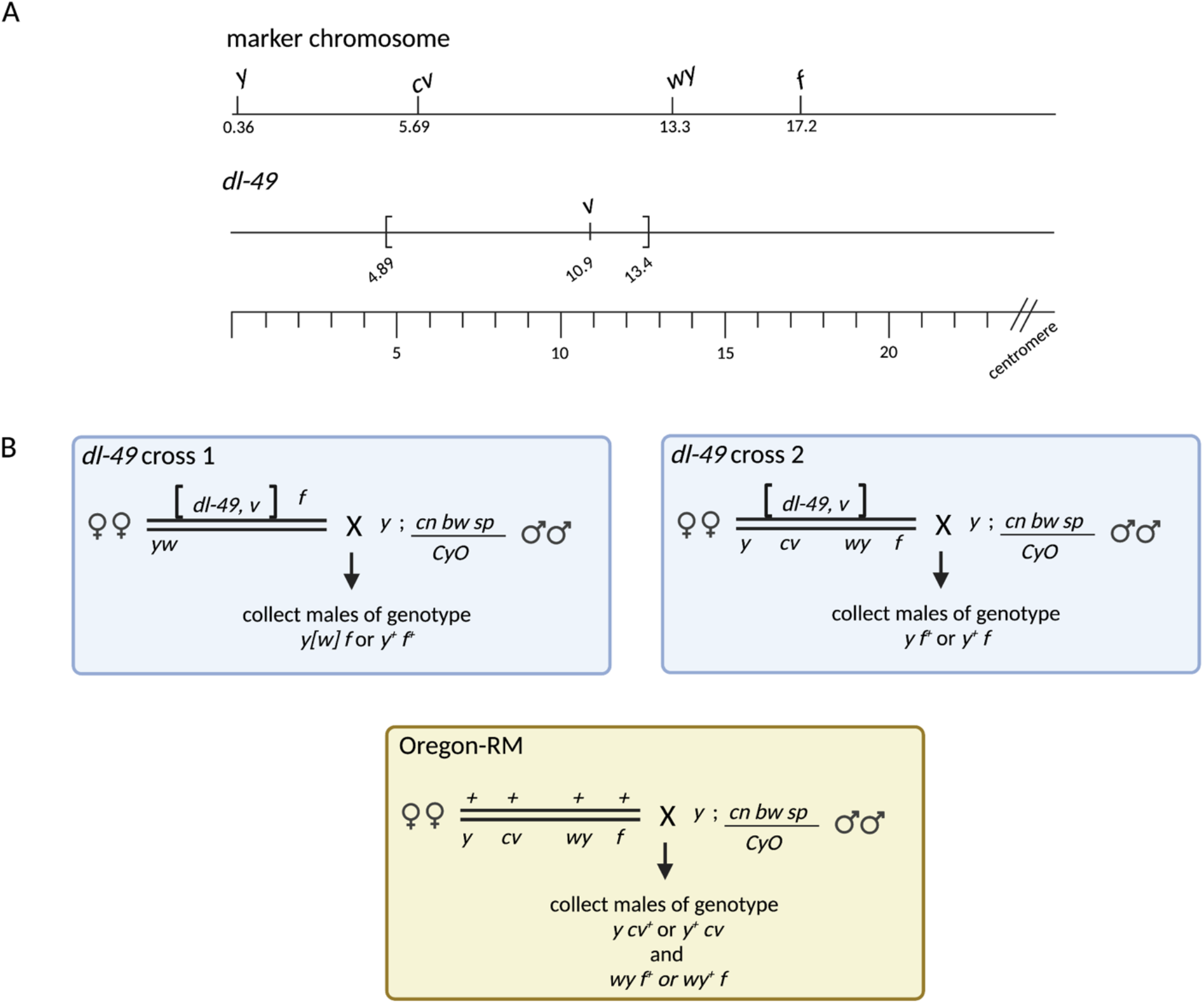
A) Genomic coordinates of the phenotypic markers and the *dl-49* inversion breakpoints. The inversion breakpoints of *dl-49* are at positions 4.89 Mb and 13.4 Mb. All numbers are in Mb using the coordinates from genome version dm6. B) Cross schemes for all three experimental set ups. COs were identified by scoring for new combinations of parental alleles.

To recover crossovers in the syntenic regions of *chrX* in the full-sibling wildtype *Oregon-RM* stock, we generated females that were *+/ y^1^ cv^1^ IP3K2^wy-74i^ f^1^* (Figure 2). These females were crossed to males of the reference genome stock *y; cn bw sp/CyO*. We collected male offspring that resulted from crossovers between *y^1^ and cv^1^* or between *IP3K2^wy-74i^* and *f*.

We were not able to directly select for noncrossover gene conversion events in any of the above crosses. To recover offspring that potentially resulted from NCOGC events, we sequenced 100 randomly chosen male offspring that maintained the parental genotype in the *dl-49* and Oregon-RM full sibling crosses.

### Illumina whole genome sequencing

We extracted genomic DNA from frozen single flies using the Qiagen DNeasy Blood and Tissue kit (Qiagen #69506) with some modifications. Single flies were homogenized using a pestle (Kimble #749520-0090) in 180 ul of ATL buffer, after which 5 ul of RNase A was added (10 mg/ml, Thermo Scientific #EN0531). We followed all other standard manufacturer protocols, except we eluted the DNA in either 40 or 50 ul of 10 mM Tris (pH 8.0) to avoid incorporating EDTA into the final DNA.

To prepare libraries for sequencing, we used the Illumina DNA Prep kit using the standard manufacturer protocols (cat no #20018705, previously known as the Nextera DNA Flex Library Prep). Libraries were amplified using 6 or 7 rounds of PCR depending on the exact concentration of the genomic DNA. We used the IDT for Illumina UD Indexes, sets A-D (#20027213, #20027214, #20027215, #200272136). The final concentration of the libraries was determined using the ThermoFisher Qubit 1X dsDNA High Sensitivity Kit (#Q33230) and a Qubit 4 Fluorometer. The 260/280 and 230/260 ratios were quantified on a Denovix DS-11 Spectrophotometer.

Libraries from experimental set up 1 were sent to Nationwide Children’s Hospital Genomics Services Laboratory and sequenced on an Illumina NovaSeq 6000 in 2 lanes with S1 2×150 chemistry. Libraries from experimental set up 2 were sent to the University of Minnesota Genomics Center and sequenced on an Illumina NovaSeq 6000 in 2 lanes with S4 2×150 chemistry. All data are available in NCBI Bioproject SUB12271178.

### Nanopore sequencing

DNA for Nanopore sequencing was prepared from 20 females of the *dl-49, v^Of^* stock generated in this study using the method in (31). Libraries for sequencing were prepared using the ligation sequencing kit (SQK-LSK110) starting with 2 ug of DNA and following the manufacturers’ instructions, except that the ligation time was extended to 30 minutes. Prepared libraries were quantified with a Qubit Fluorometer (ThermoFisher) and 600 ng of prepared library was loaded onto a R9.4.1 PromethION flow cell that had been previously run for 48 hours and run for 24 hours. FASTQ files were generated using Guppy version 6.1.1. (Oxford Nanopore Technologies) and aligned to *Drosophila melanogaster* genome version dm6 using minimap2 (32). Sniffles2 was used to identify structural variants (33).

### Analysis to identify COs and NCOGCs

Illumina short read data was aligned to the *Drosophila melanogaster* genome (dm6) using minimap2 (32). Single nucleotide variants (SNVs) were identified using samtools (34). COs and NCOGCs were identified as previously described (27, 35). Briefly, for all parental stocks, SNVs on the X chromosome that were unique to one of the two parental lines were identified. For each offspring, the parental origin of each SNV was determined and any SNV that represented a switch from one parent to the other was flagged as a possible CO or NCOGC event.

### Statistical analyses

When analyzing CO frequencies in *dl-49* heterozygotes, we limited our analyses to COs that occurred between the proximal breakpoint and *f*. COs that occurred proximal to *f* were recovered in the nonrecombinant offspring and had not been selected for, thus we excluded them from our analyses (Figure 3). Because we only sequenced a subset of the total COs recovered, we did not measure CO frequencies using centiMorgans, the traditional measurement of CO frequency. Instead, we normalized CO frequencies within each interval by the total number of COs that were sequenced for each genotype. This provided a more accurate representation of the distribution of COs within our samples.

**Figure 3.**
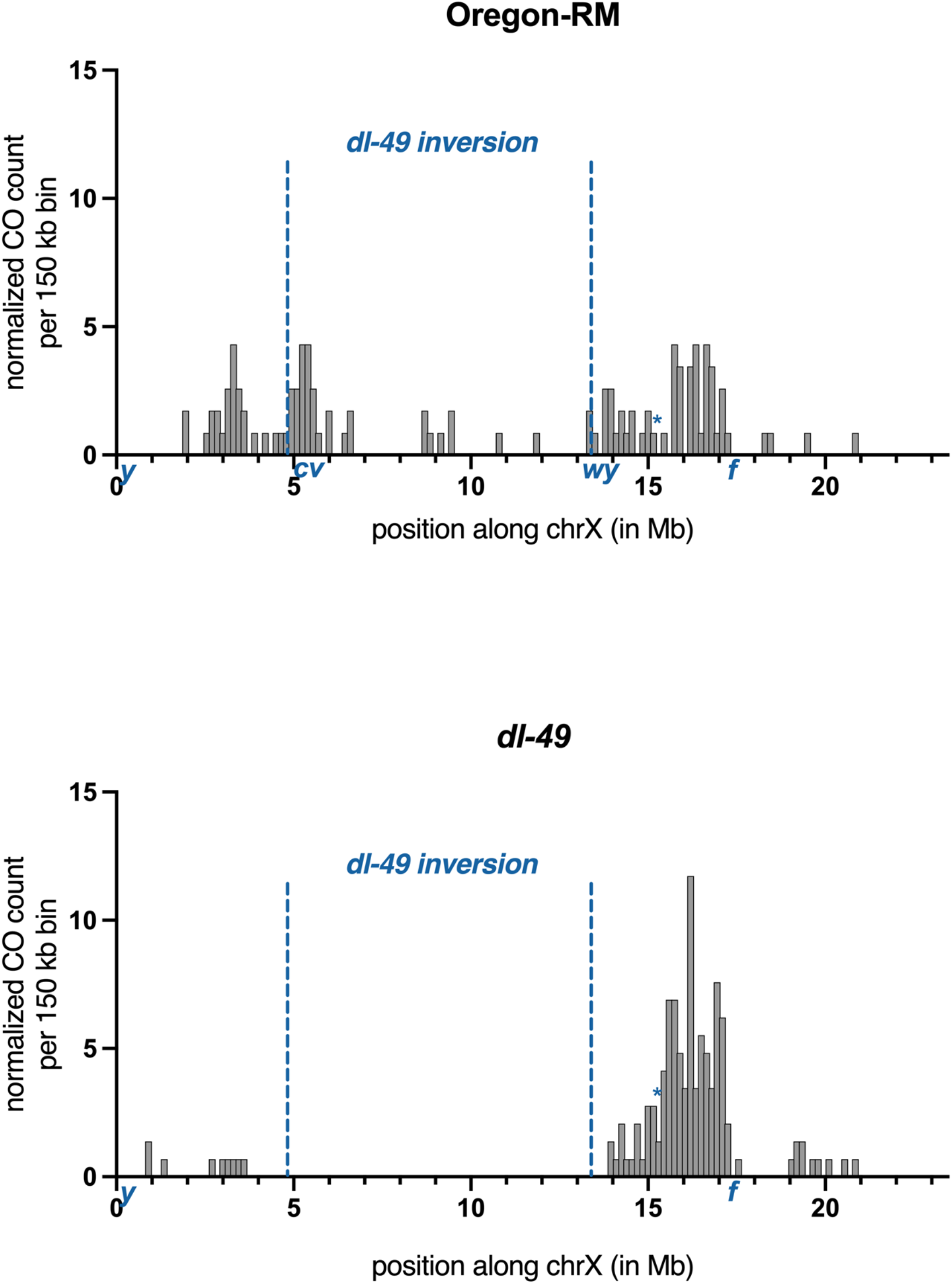
Location and frequency of COs recovered in Oregon-RM and *dl-49* heterozygotes. COs that occur between *cv* and *wy* were from samples that had more than one CO on the chromosome. These COs were not included in downstream analyses. Asterisk shows the location of the 15.375 interval. CO counts were normalized to sample size by dividing the number of COs per interval by the total number of COs. In Oregon-RM, CO distribution between the *dl-49* breakpoint and *f* are not correlated with distance (Spearman’s rank correlation, r = 0.18, p = 0.35). In *dl-49* heterozygotes, CO distribution between the *dl-49* breakpoint and *f* are correlated with distance (Spearman’s rank correlation, r = 0.47, p = 0.01).

When analyzing CO frequencies in Oregon-RM, we selected for COs that occurred between *y* and *cv* or *IP3K2^wy-74i^* and *f*. However, 21 of the offspring we sequenced had two COs on the same chromosome, one of which occurred between *cv* and *IP3K2^wy-74i^*. These COs are shown in Figure 3, but were not included in any analyses.

All Fisher’s exact tests were performed using QuickCalcs at https://www.graphpad.com/quickcalcs/. All other statistical tests were performed in GraphPad Prism version 9.4.

## Results

### Identification of COs in wildtype and dl-49

To determine the mechanism of how heterozygous inversions suppress COs in the regions outside of the breakpoints, we built fine-scale genetic maps of rare CO and NCOGC events within approximately 4 Mb of the breakpoints. We used the X chromosome inversion *dl-49* because it is completely euchromatic *(28)* and visible recessive markers exist at convenient locations to facilitate identification of COs (Figure 2). We performed this experiment in two different genetic backgrounds. First, we generated females that were heterozygous for *dl-49, v, f* and a *yw* marker chromosome, then recovered COs that occurred between *y* and *f* (Figure 2). The map length between *y* and *f* was 1.76 cM in this genetic background, confirming that the *dl-49* inversion does suppress CO formation (Table 1). Second, in order to have a directly comparable wildtype dataset from the same genetic background, we created congenic *dl-49* and Oregon-RM stocks by out crossing the *dl-49* inversion into the Oregon-RM background for nine generations and creating full-sibling stocks at the tenth generation (Methods and Supplemental Figure 2). We generated females that were heterozygous for *dl-49, v* and a *y cv wy f* marker chromosome, then recovered COs between *y* and *f* (Figure 2). The map length between *y* and *f* was 2.01 cM in this genetic background, however, we were unable to discriminate between a CO and nondisjunction in this cross, so this is an overestimate of map length (Methods and Table 1). Lastly, we generated females using the full-sibling Oregon-RM stock that were heterozygous for the *y cv wy f* marker chromosome and recovered COs between *y* and *cv* or *wy* and *f* in order to limit our analyses to the syntenic regions of the inversion breakpoints (Figure 2). The map length was 9.5 cM for *y-cv* and 10.4 for *wy-f* (Table 1). These map lengths are statistically different (Fisher’s exact test, p < 0.0001) confirming that *dl-49* suppresses crossovers to approximately 10% of wildtype levels in the *y-f* interval.

**Table 1.**
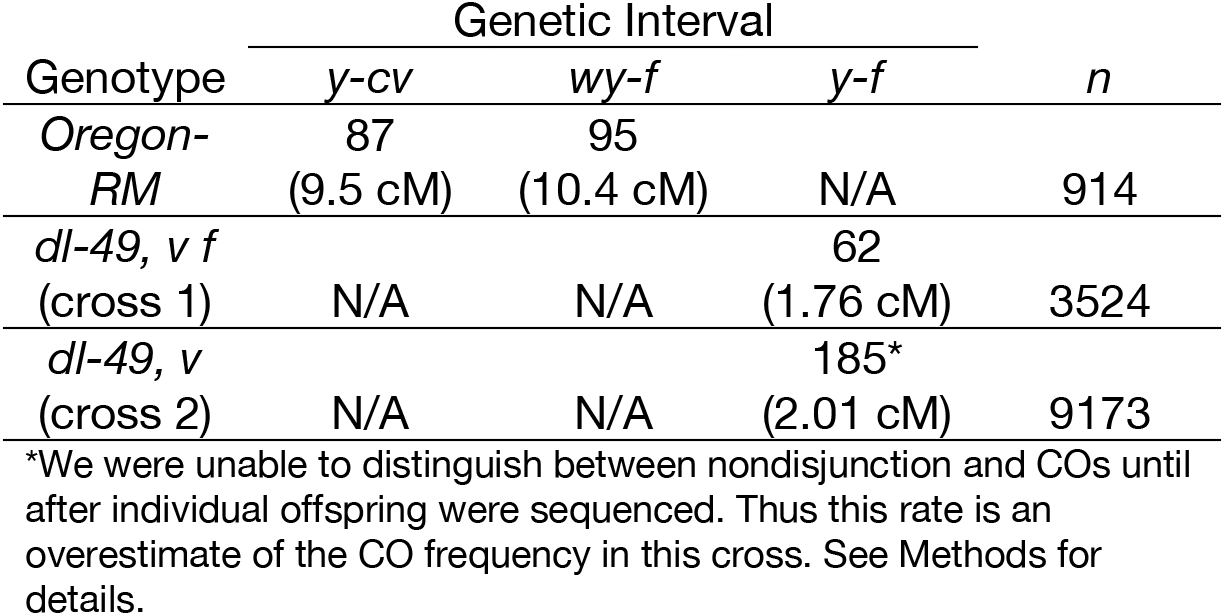
Crossover frequencies in the specified genetic intervals for the full-sibling wildtype Oregon-RM and the *dl-49* inversion. Number of recovered offspring are shown, and the corresponding cM is shown in parentheses below.

We next identified the precise location of 145 COs that occurred between *y* and *f* in *dl-49* heterozygotes and 96 COs between *y* and *cv* or *wy* and *f* in Oregon-RM (Supplemental File 1). We randomly selected individual recombinant male offspring from each genotype and used whole genome Illumina sequencing to determine the exact location of the CO by identifying where the genotype switched from the *dl-49* or Oregon-RM genotype to the marker chromosome genotype (Figure 2). The *dl-49* and *y cv wy f* stocks had a distribution of approximately 1 SNV every 367 bp on *chrX* and the *Oregon-RM* and *y cv wy f* stocks had a distribution of approximately 1 SNV every 346 bp on *chrX*. These SNV densities allowed us to pinpoint the location of the CO with very high resolution. We analyzed CO frequencies by normalizing for sample size and binning CO counts into 150 kb non-overlapping windows (Figure 3 and Supplemental Figure 4). COs from both *dl-49* crosses were combined into one dataset for further analysis because the CO distributions were not statistically different (Supplemental Figure 3, Kolmogorov-Smirnov test, p = 0.61).

### Distribution of COs in wildtype and dl-49

Because only nine of the 145 COs in *dl-49* heterozygotes occurred between *y* and the distal inversion breakpoint, so we wondered if a cryptic inversion polymorphism could be suppressing COs in this region. We performed whole-genome long-read sequencing of *dl-49* on the Nanopore platform followed by *de novo* assembly and did not identify any additional structural variants. Previous studies of the *dl-49* chromosome performed in different decades in different labs show that COs are reduced to nearly 0% in this region, so this effect is not unique to the current experiment [11, 12, 18]. Since there were only nine COs on the distal end, we restricted our analyses to COs on the proximal end.

We next asked if CO frequency is correlated with distance from the proximal inversion breakpoint. After normalizing for sample size, we found that in Oregon-RM, CO frequencies between position 13.4 Mb (the position of the *dl-49* breakpoint) and *f* are not correlated with distance (Spearman’s rank correlation, r = 0.18, p = 0.35). However, in *dl-49* heterozygotes, CO frequencies between the inversion breakpoint and *f* are correlated with distance (Spearman’s rank correlation, r = 0.47, p = 0.01).

After visual inspection, we noticed a clear change in CO frequencies at position 15.375 Mb in the *dl-49* heterozygotes. CO frequencies between the inversion breakpoint and 15.375 Mb were always below 4 COs per interval, while CO frequencies between 15.375 Mb and *f* were always at least 5 COs per window (Figure 3 and Supplemental Figure 4, the only exception to this was the interval at 17.55 Mb). We divided the COs into two windows based on this distribution and found that the proportion of COs distributed into these two large windows is statistically different in *dl-49* heterozygotes than in Oregon-RM, with fewer COs within 2 Mb and more COs within 2-4 Mb (Fisher’s exact test, p = 0.026, Figure 3). This suggests that inversion breakpoints have the strongest inhibitory effect on CO frequency over a distance of 2 Mb, while COs 2-4 Mb from the breakpoint increase in frequency. Importantly, the bimodal effect of the inversion breakpoint is no longer statistically significant if the boundary is moved to position 15.45 Mb (Fisher’s exact test, p = 0.12), consistent with the existence of a defined position after which the effect of the inversion breakpoint changes.

There was a notable absence of COs within approximately 500 kb of the proximal inversion breakpoint, with the closest being 470 kb outside the inversion breakpoint (Figure 3 and Supplemental File 1). In Oregon-RM, there were 5 COs in this same interval (Figure 3 and Supplemental File 1). The difference in CO frequency in this 500 kb window is statistically different (Fisher’s exact test, p = 0.0085). This is consistent with a model where COs are completely suppressed within approximately 500 kb of the inversion breakpoint, although we cannot rule out that CO frequencies are so low in this region that we simply did not recover them given our sample size.

### Distribution of NCOGCs in wildtype and dl-49

It remains unclear how NCOGCs are affected by inversion breakpoints because their small size makes them difficult to detect and the existing data sets are small. While we could not select for NCOGCs in our system, we reasoned that by sequencing enough nonrecombinant offspring, we would occasionally recover NCOGCs that occurred close to the inversion breakpoints. We performed whole-genome Illumina sequencing on 100 randomly selected nonrecombinant male offspring from *dl-49* heterozygotes and Oregon-RM females to identify NCOGCs. These nonrecombinant offspring were selected from the same crosses used to select for the COs described above.

We recovered a total of 51 unique NCOGCs from *dl-49* heterozygotes and 23 unique NCOGCs from Oregon-RM across the entire X chromosome (Figure 4 and Supplemental File 1). While this difference in frequency is statistically significant (Fisher’s exact test, p < 0.0001), it is likely that the increase in NCOGC frequency in *dl-49* heterozygotes is at least partially due to transmission distortion; only chromosomes without a CO within the inversion are transmitted to the offspring, which selects for chromosomes that had NCOGCs instead. Regardless, the net effect is that NCOGCs are transmitted at a higher rate than in wildtype, similar to previous observations using multiply inverted balancer chromosomes (27)

**Figure 4.**
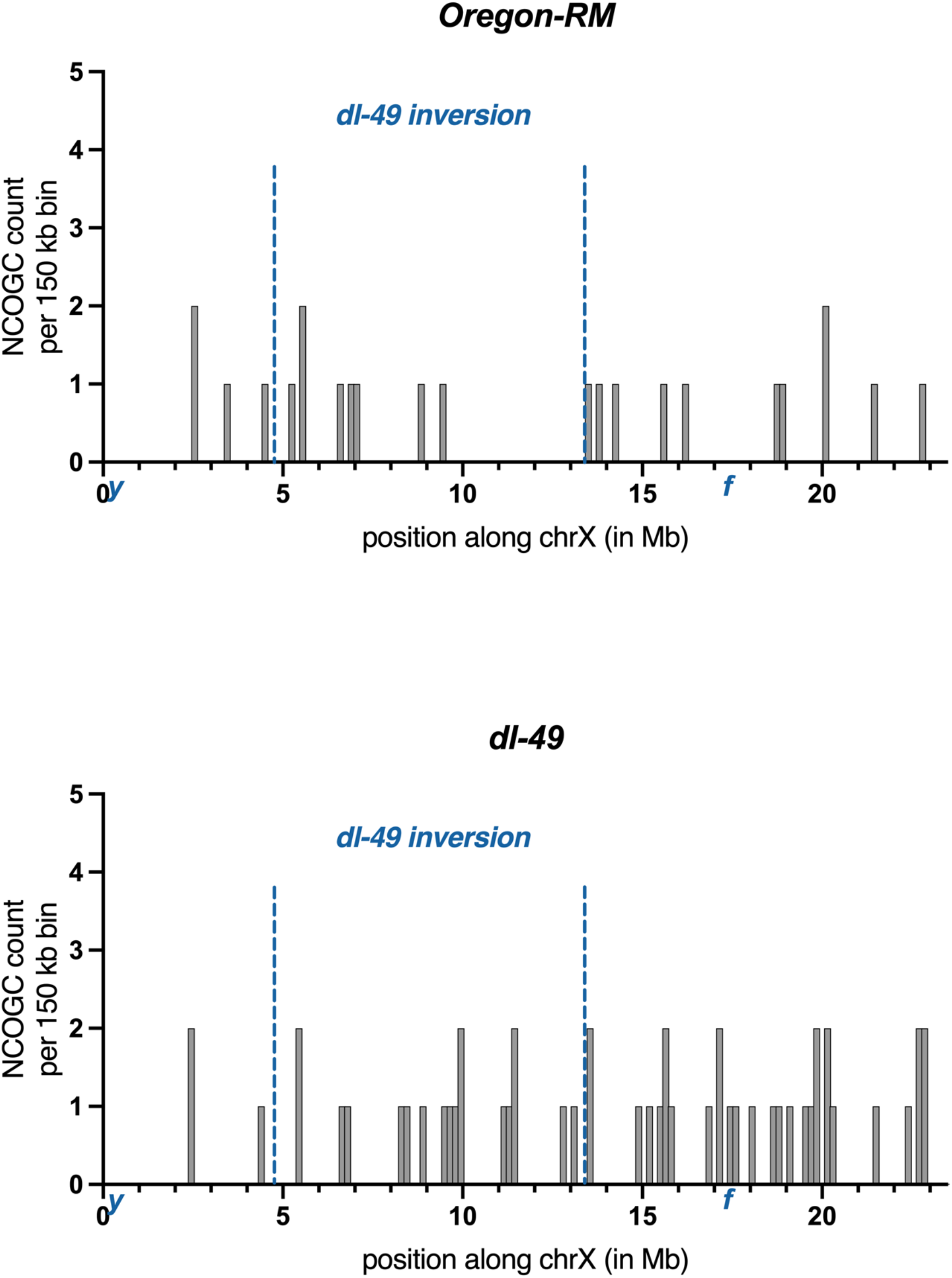
Location and frequencies of NCOGCs recovered from Oregon-RM and *dl-49* heterozygotes. Frequencies are raw counts per 150 kb window.

Only three of the 51 NCOGCs in *dl-49* heterozygotes occurred between the telomere and the distal *dl-49* inversion breakpoint, whereas four of the 23 NCOGCs in Oregon-RM occurred in this region (Figure 4 and Supplemental File 1). We re-examined the local SNV density in this region and found only 1 SNV every 81,540 bp. NCOGC length is approximately 400 bp in *D. melanogaster* (24), thus we did not have enough SNV density to detect NCOGCs in this area and excluded it from further analysis.

In the *dl-49* heterozygotes, 30 of the NCOGCs occurred between the proximal breakpoint and the centromere, and two of these 30 NCOGCs were within 500 kb of the inversion breakpoint (Figure 4 and Supplemental File 1). In the same region of Oregon RM, there were 11 NCOGCs, 2 of which were within the same 500 kb interval (Figure 4 and Supplemental File 1). These proportions are not statistically different (Fisher’s exact test, p = 0.29), suggesting that NCOGCs occur in this 500 kb region at the same frequency in Oregon-RM and *dl-49* heterozygotes. Measurements of NCOGC frequencies will always be underestimates, but because our experimental set up allows us to directly compare the *dl-49* heterozygotes to Oregon-RM, our data show that NCOGCs occur near inversion breakpoints at the same frequency as in wildtype. This also suggests that the genomic regions near inversion breakpoints are competent to form DSBs.

### Pre-meiotic or somatic NCOGCs

There were 5 instances of NCOGCs that occurred in identical locations in different individuals from the *dl-49* heterozygotes (Supplemental File 1). Since DSBs do not form in hotspots in Drosophila species (36), it is most likely that these NCOGCs occurred pre-meiotically in the germline. We did not recover any similar instances in Oregon-RM. Interestingly, other reports of similar pre-meiotic jackpot events occurred in genotypes with at least one heterozygous inversion, suggesting that they may affect recombination or other DNA repair mechanisms in non-meiotic cells (23, 26).

## Discussion

It has been known for almost 100 years that heterozygous inversions suppress COs in the regions immediately outside of the breakpoints (1, 11, 12), however the underlying mechanism was unknown. Based on the data described here, we propose the following model. The genomic regions near inversion breakpoints are able to form DSBs at wildtype frequencies. Within approximately 500 kb of the inversion breakpoint, DSBs are preferentially repaired into NCOGCs. Between 500 kb and 2 Mb, DSBs can be repaired into COs but this repair outcome happens at a much lower frequency than wildtype, leading to an overall reduced CO frequency in these regions. Between 2-4 Mb, CO frequencies increase above wildtype for unknown reasons. While there is a dramatic shift in the distribution of COs outside of inversion breakpoints, there is still a net decrease in CO frequencies.

Importantly, our data argue that the decrease in CO frequency is not caused by a decrease in DSB formation. The frequency and location of the NCOGCs clearly show that DSBs can occur in genomic regions very close to the breakpoints. We and others have previously shown that the interchromosomal effect – the genome-wide increase in CO frequency caused by a heterozygous inversion – is also not mediated by DSB number (27, 37); rather, the number of DSBs stays the same and the increased COs form at the expense of NCOGCs (27). Together, these data suggest that, in *D. melanogaster*, plasticity in local and global CO frequencies due to heterozygous inversions is generally mediated by altering recombination repair outcome as opposed to changes in DSB number.

We also show that CO suppression is absolute within 500 kb and strong within 2 Mb of the inversion breakpoint. There is remarkable consistency between our direct measurements of CO frequencies and those obtained in other contexts. Indirect measurements of CO frequencies at the distal end of *TM3*, a multiply inverted third chromosome balancer, suggest that COs can occur as close as 2 Mb to an inversion breakpoint (35). Direct measurements of CO frequencies during the interchromosomal effect identified COs as close as 1.7 Mb to an inversion breakpoint (27). Direct measurements in interspecies crosses of *D. pseudoobscura* and *D. persimilis* show that COs are suppressed up to 2-3 Mb from the breakpoint (19, 20). Interestingly, in *Arabidopsis thaliana*, COs are only suppressed approximately 10 kb outside of inversion breakpoints, with some occurring as close as 1 kb (38); this intriguing difference could be explained by the different relationship between DSB formation and chromosome synapsis in Arabidopsis and Drosophila species (39, 40).

Previous data left uncertainty about rates of NCOGCs near inversion breakpoints. For example, our previous work showed that NCOGCs can occur within 500 kb of the breakpoints on multiply inverted balancer chromosomes (27). However, without a wildtype dataset to compare to, it was unclear whether NCOGCs in these regions were occurring at wildtype or reduced frequencies. Similarly, previous work in *D. pseudoobscura* found that of 32 unique NCOGCs, only one was within 500 kb, although the sample size in that study may have been simply too small to detect NCOGCs near inversion boundaries. In the current work, we clearly show that NCOGC frequencies in the regions outside inversion breakpoints are statistically the same in the syntenic regions in wildtype (Figure 4).

### Crossover suppression at the distal end of dl-49

We observed a striking paucity of COs on the distal end of *dl-49*. COs are generally reduced in sub-telomeric regions and peri-centromeric regions (24). *dl-49* is only 4.89 Mb from the telomere but is approximately 10 Mb away from the peri-centromeric heterochromatin. We suggest that the combined effects of the telomere and the inversion breakpoint severely restrict CO formation on the distal end, but the proximal breakpoint is too far away from the centromere to be affected. Consistent with this idea is the behavior of the *ClB* inversion, a larger X chromosome inversion with breakpoints at cytological position 4A5 (approximately position 4.1 Mb) and 17A6 (approximately position 18.35 Mb) [36]. Similar to *dl-49*, COs between *y* and the distal breakpoint are very low (0.77 cM), but unlike *dl-49*, COs are severely suppressed between the proximal breakpoint and the centromere [37]. The proximal breakpoint is only 5 Mb from the pericentromeric chromatin, consistent with the idea that in *ClB*, the centromere effect combined with the inversion breakpoint severely limits CO formation.

Alternatively, the lack of COs on the distal end of *dl-49* could be explained by a pairing defect. The 4.89 Mb between the distal breakpoint and the telomere may not be sufficient for chromatin to maintain stable pairing. Lastly, DSB frequency might be reduced in this region for unknown reasons. Resolving the cause for this interesting observation would require the recovery of all NCOGCs in this region, but unfortunately the poor SNV density prevents us from knowing the true NCOGC frequency.

### Chromosome pairing and synaptonemal complex formation in structural variant heterozygotes

Because the regions near heterozygous inversion breakpoints are clearly competent to form DSBs, the next obvious question is why do COs form at such low levels in these intervals? The synaptonemal complex (SC) – a tripartite proteinaceous structure that forms between homologous chromosomes during meiosis – is thought to regulate CO formation (41). Both the chromosome axes and the SC are likely under increased physical stress near inversion breakpoints due to formation of inversion loops (Figure 1).

Surprisingly, there are no overt chromosome pairing or SC defects in inversion heterozygotes. Analyses of the SC protein C(3)G show that SC does form along the length of a heterozygous *FM7* balancer. Additionally, the heterozygous *FM7* balancer is paired with its non-inverted homolog in 70-80% of meiotic nuclei, depending on the location of the FISH probe (37). In translocation heterozygotes, which also suppress COs, the SC is built along the length of the chromosome, although it appears to be missing at the translocation breakpoint in 10-20% of nuclei (42). Translocation heterozygotes are similarly paired in 80-90% of meiotic nuclei (42). While the severity of the CO suppression in structural variant heterozygotes does not seem to correlate well with the mild pairing and SC defects, it is possible that there are defects in SC formation that cannot be detected with simple immunostaining.

We are left with trying to understand how heterozygous inversion breakpoints suppress CO formation even though DSBs are made, NCOGCs form at wildtype levels, chromosomes are mostly paired, and the SC is at least superficially intact. One possible explanation is that increased physical stress at the inversion breakpoints prevents DSBs from being repaired into COs very locally; as the distance from the breakpoint increases, the stress dissipates, and normal SC structure allows DSBs to be repaired into COs. Indeed, Sturtevant proposed that defects in chromosome synapsis could be a reason COs were suppressed by inversions when he first observed this in 1926 (1). Others have proposed that inversion breakpoints might act as CO interference signals (37), which also dissipate with distance (43), or that COs require long uninterrupted blocks of synapsis in order to occur (42). Our data are certainly consistent with these ideas and cannot discriminate between them.

It is possible that there is hyperlocal asynapsis at inversion breakpoints that could not be detected with previous imaging approaches; however, if this were the case, we would not have been able to detect NCOGCs. Clearly, regions near the breakpoints are able to interact with the homolog at least long enough to form NCOGCs. It is possible that unstable interhomolog interactions in these regions (due to inversion loops) prevent recombination intermediates from being repaired into COs and that the NCOGCs are representative of short-lived interhomolog interactions. Given this, we favor a model where heterozygous inversion breakpoints destabilize interhomolog interactions locally, such that chromosome interactions are stable enough to facilitate DSB formation and recombination in general, but not CO formation. Future experiments detailing how inversion breakpoints impact recombination mechanics at the molecular level and super-resolution imaging analysis of the SC should provide insight into this issue.

## Acknowledgements

We would like to thank Nadia Singh for helpful advice on experimental set up and statistical analyses and R. Scott Hawley for the isogenized Oregon-RM stock. We would also like to thank members of the Crown lab, Rob Ward, Helen Salz, and Talia Hatkevich for helpful comments on the manuscript and Angie Miller for help in preparing figures. This work was supported by R35 GM137834 to KNC.

## Author Contributions

HL, EB, SH, and MG were responsible for investigation. DEM was responsible for formal analysis and writing. KNC was responsible for investigation, conceptualization, and writing.

## Supplemental Figures

**Supplemental Figure 1.**
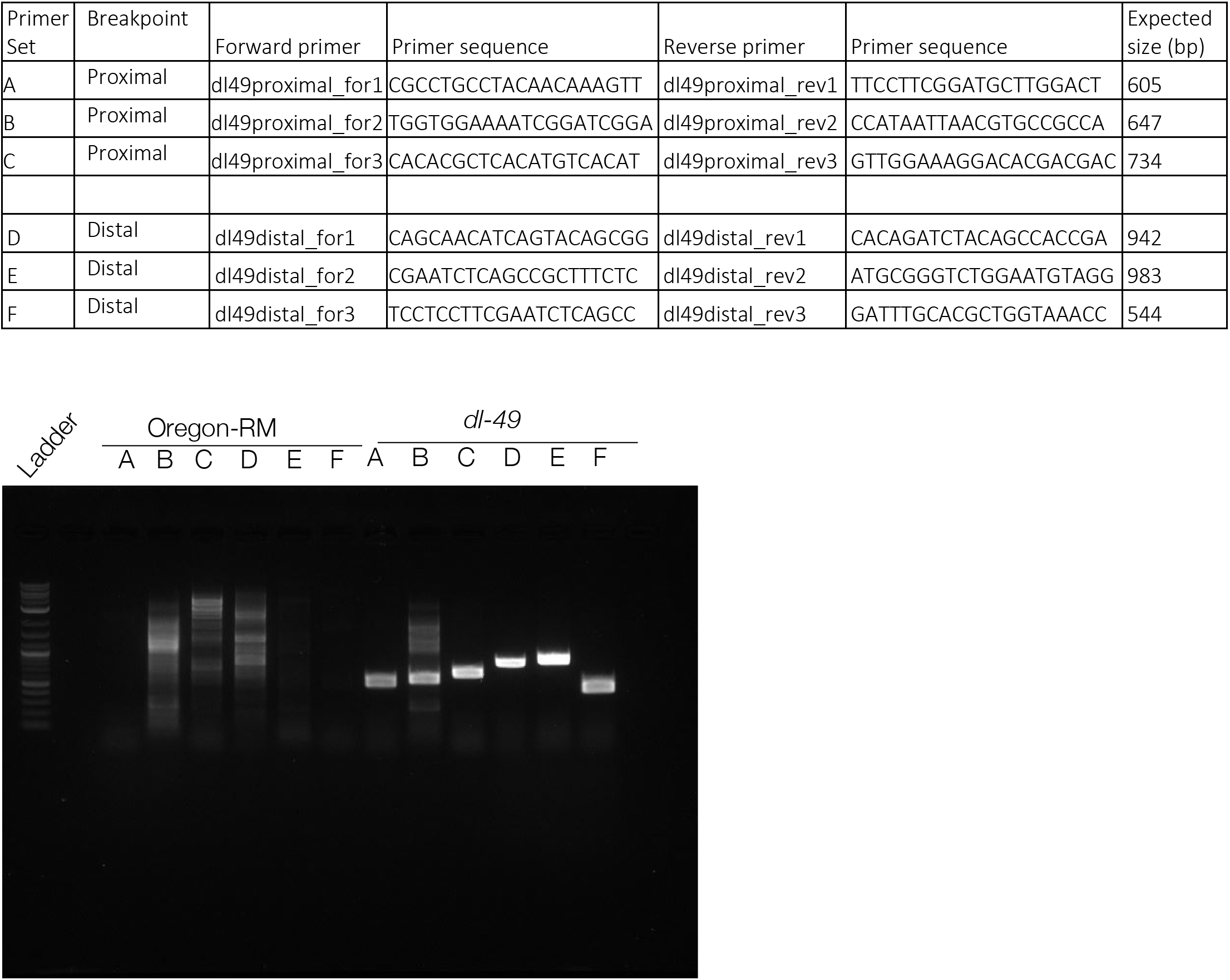
PCR primers used to detect the *dl-49* breakpoints. PCR results show that primer sets fail to amplify a specific band in Oregon-RM, but do amplify specific bands of the expected sizes in *dl-49*, confirming that the expected breakpoints are present.

**Supplemental Figure 2.**
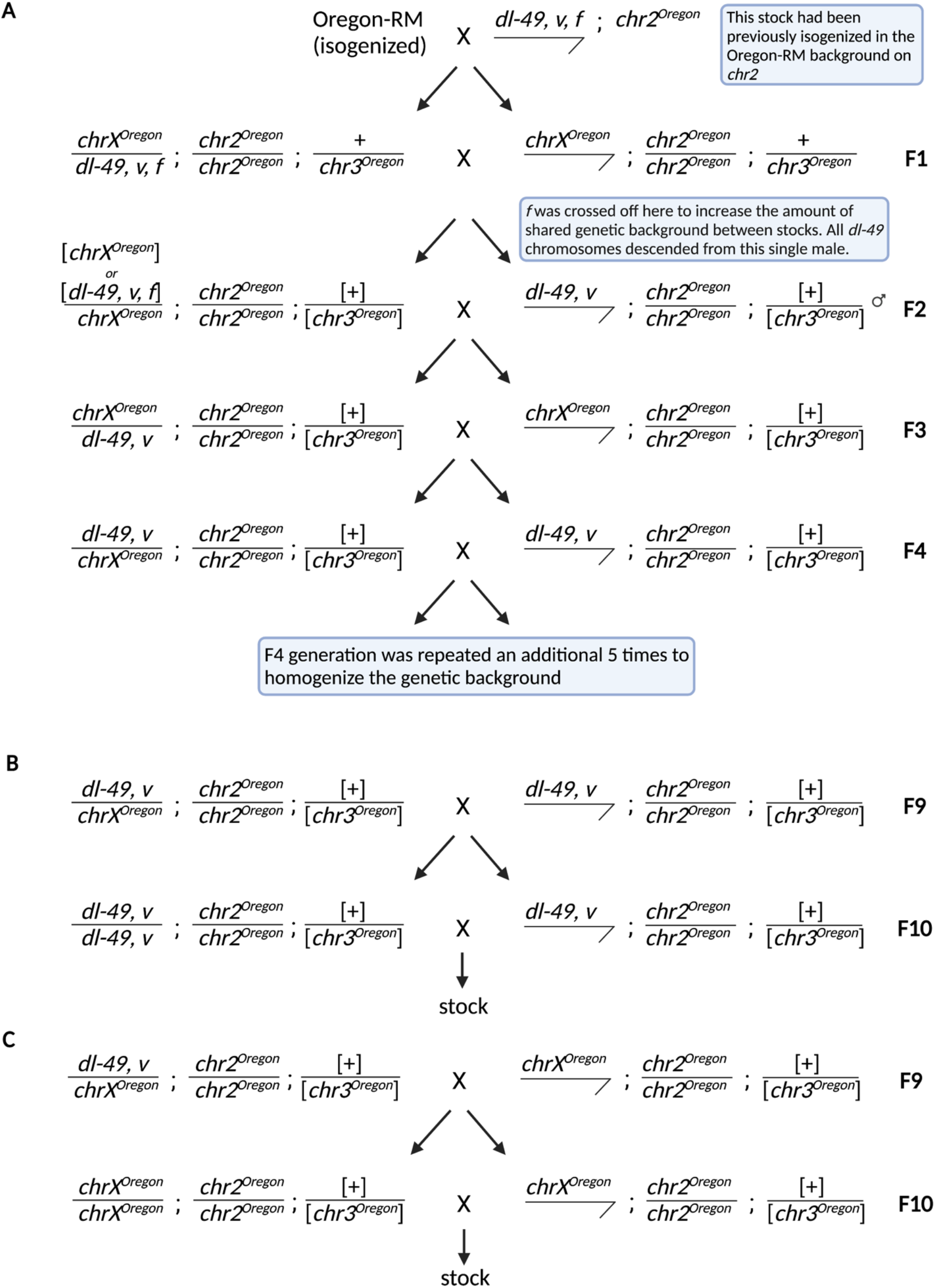
Crosses used to generate full sibling Oregon-RM and *dl-49* stocks. A) We previously isogenized *dl-49* and crossed in an isogenized *chr2* from Oregon-RM. This stock was used for cross 1 in Figure 2. The next several generations were needed to create a stock that was heterozygous for *dl-49* and *chrX* from Oregon-RM and that shared genetic background on *chr3*. Full-sibling stocks were created at generation 10 by crossing *dl-49* heterozygotes to either *dl-49* males (B) or Oregon-RM males (C). This cross scheme did not result in isogenized 3^rd^ chromosome, but did create a shared genetic background.

**Supplemental Figure 3.**
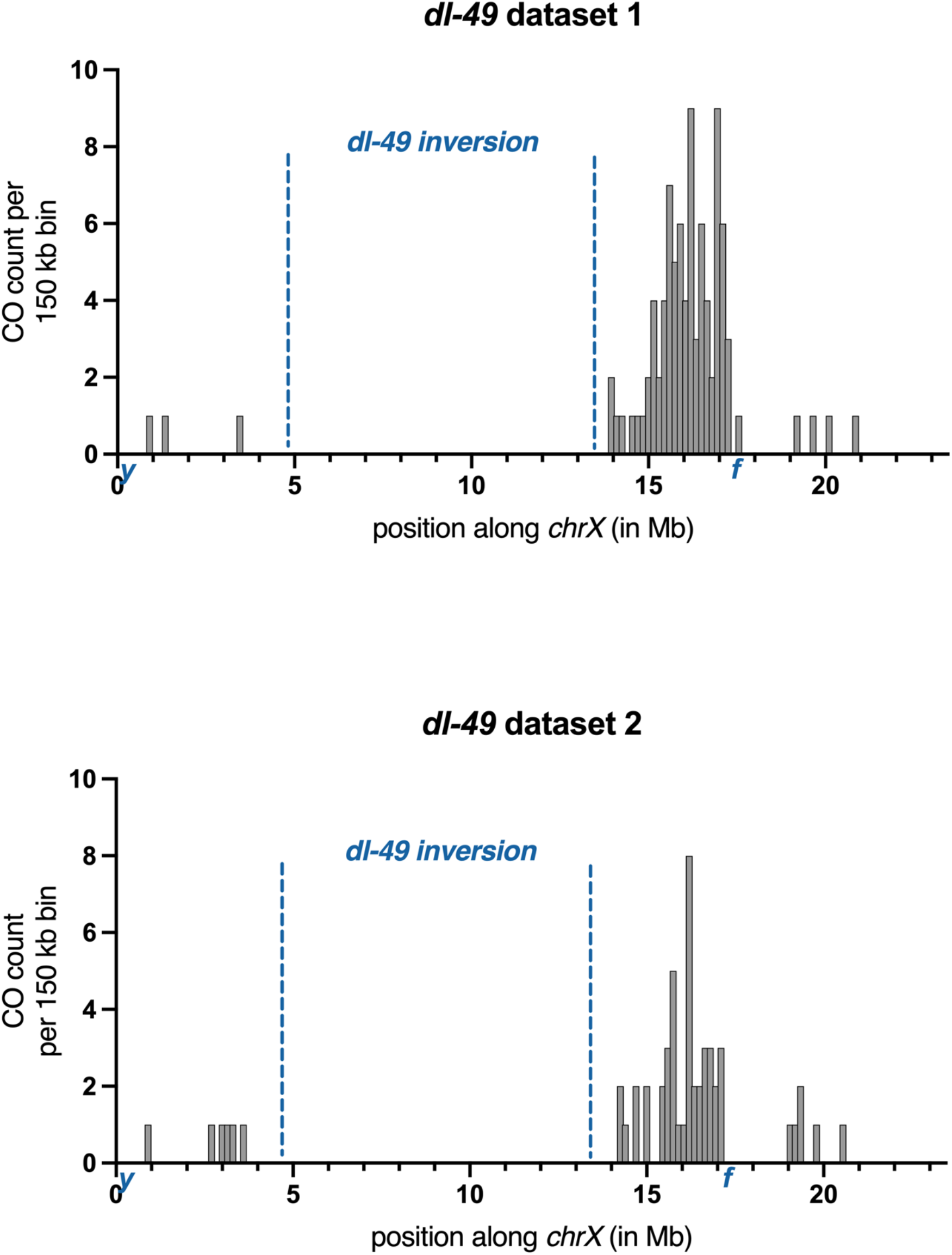
CO frequencies in *dl-49* heterozygotes from cross 1 and cross 2 (Figure 2). These distributions are not significantly different from each other and were combined into one dataset (Kolmogorov-Smirnov test, p = 0.61).

**Supplemental Figure 4.**
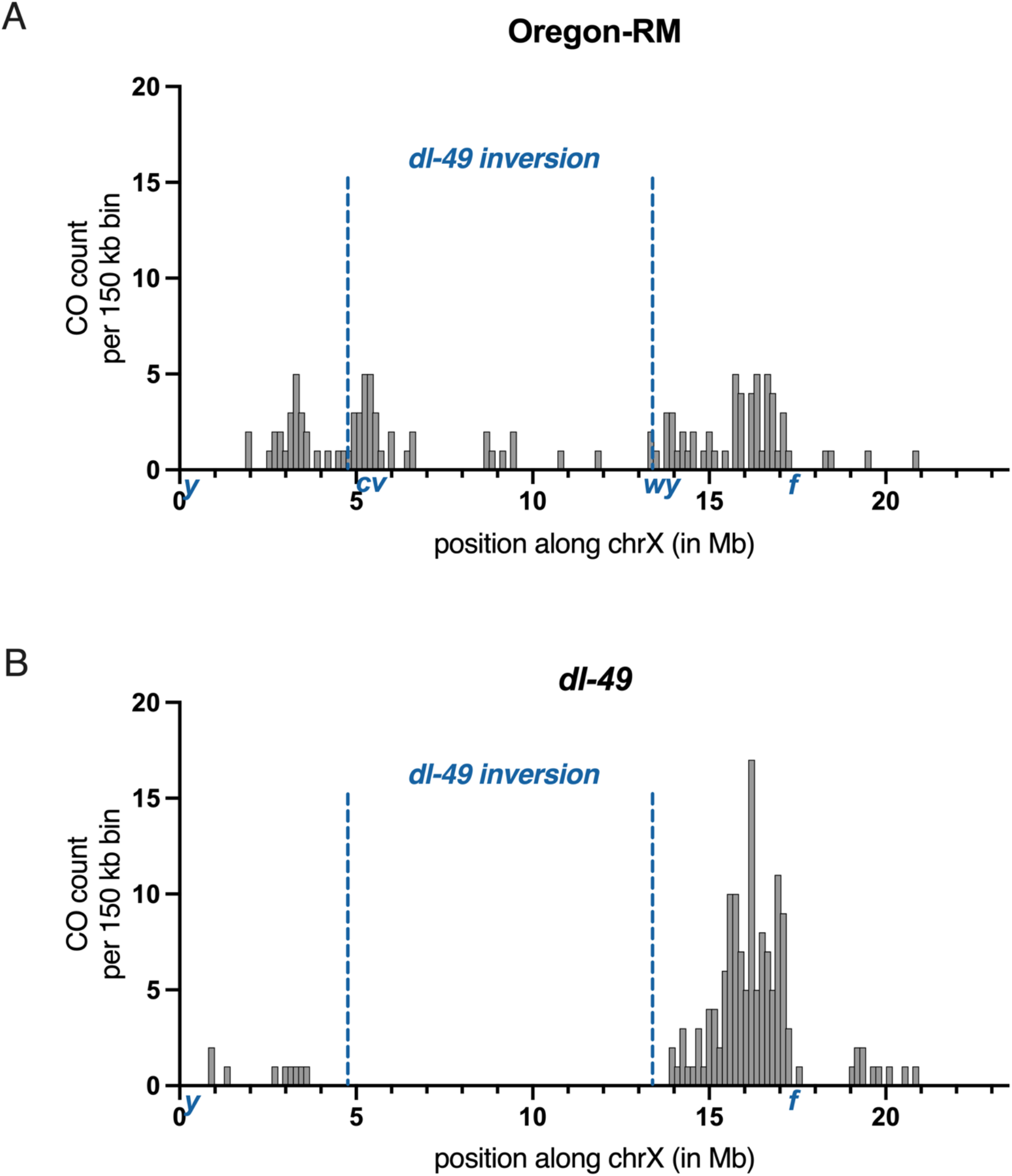
A) Raw counts of CO frequencies in Oregon-RM. 96 COs between *y* and *cv* or *wy* and *f* were sequenced. COs that occur between *cv* and *wy* were from samples that had more than one CO on the chromosome. These COs were not included in any analysis. B) Raw counts of CO frequencies from *dl-49* heterozygotes. 145 COs between *y* and *f* were sequenced.

Supplemental File 1

See attached spreadsheet

